# Coupling Anaerobic–Oxic and Integrated Fixed–Activated Sludge Processes for Removal and Transformation of Refractory Organic Matter in Livestock and Poultry Digestate

**DOI:** 10.1101/2025.04.15.649029

**Authors:** Bingxin Li, Heyang Yuan, Zheng Ge

**Author notes:** Corresponding authors. E–mail. Intended for Journal Name: ***Bioresource Technology***. Type of contribution: ***Research Article***.

## Abstract

Effective treatment of livestock and poultry digestate remains challenging due to its high content of refractory organic matter. In this study, a coupled anaerobic–oxic (A/O) and integrated fixed–film activated sludge (IFAS) system was investigated to evaluate its capacity for organic removal and molecular transformation. The IFAS system outperformed the A/O system, achieving an organic removal efficiency of 43%, compared to 17% in the A/O process. Molecular analyses revealed distinct transformation pathways: the A/O process tended to increase the molecular weight and structural complexity of dissolved organic matter, whereas the IFAS process promoted oxidative fragmentation, resulting in the formation of smaller, oxygen–rich compounds. These changes were reflected in key molecular indices (DBE, NOSC, AI_mod_) and the accumulation of carboxyl–rich intermediates. Microbial community analysis associated *Bacteroidota* with the degradation of oxygenated compounds via hydrolytic processes, while *Actinobacteria* and *Deinococcota* were linked to the transformation of lignin– derived aromatics. The integration of advanced molecular characterization with microbial profiling highlights the IFAS system’s enhanced capability for both structural simplification and biodegradation of complex organics, providing a promising strategy for the treatment of high–strength organic wastewater.

## 1. Introduction

Livestock and poultry farming generates approximately 24 billion tons of wastewater annually (Koneswaran and Nierenberg, 2008), and this number is expected to exceed 100 billion tons by 2050 (Koneswaran and Nierenberg, 2008; Sigsgaard and Balmes, 2017). One of the common practices is to treat these waste streams with anaerobic digestion (Lim and Kim, 2015; Abdel-Raouf et al., 2012; Cheng et al., 2020; Yao et al., 2020), but the resulting digestate is enriched with refractory organic matter, such as phenols, aromatic hydrocarbons, and heterocyclic compounds, etc. (Chen., et al., 2023; Park et al., 2022). These substances are chemically stable and may possess antimicrobial properties that inhibit the treatment of livestock and poultry digestate (Jaeger et al., 2001; Kuhn and Suflita, 1989).

Chemical methods, such as advanced oxidation processes, are commonly applied to treat livestock and poultry digestate (Wang et al., 2016). Although this process effective in removing refractory organic matter, it involves substantial chemical dosage and operational costs (Deng and Zhao, 2015). Additionally, the strong oxidants used can interact with digestate components, producing harmful by–products and leading to secondary pollution (Domingues et al., 2021; He et al., 2009). Other chemical treatments like electrolysis and adsorption have also been investigated (Chen et al., 2023). However, electrolysis is constrained by expensive electrodes and low efficiency, limiting its scalability (Rajkumar and Palanivelu, 2004). Similarly, adsorption is limited by the cost for adsorbent regeneration and reduced capacity after reuse (An et al., 2024; Satyam and Patra, 2024). These drawbacks highlight the urgent need for more effective, cost–efficient, and scalable approaches to remove refractory organic matter from livestock and poultry digestate.

Biological treatment presents a promising and sustainable solution for the removal of refractory organic matter. Among these, the anaerobic–oxic (A/O) process has been applied to treat wastewater containing refractory organic matter (Cheng et al., 2020). For example, Yuan et al. (2017) developed an A/O process to treat mature landfill leachate and achieved 15% total organic carbon removal, with the anaerobic and oxic stages contributing 5% and 10%, respectively. The process also transformed compounds with low oxygen/carbon (O/C) ratios (0.1–0.3) into those with higher ratios (0.5–0.7). Similarly, Tian et al. (2022) reported 25% chemical oxygen demand (COD) removal from aged landfill leachate by biological treatment, with 6% and 2% removal in the anaerobic and oxic stages. Another study found that a two–stage A/O process reduced the average molecular mass of organic matter by 25%, producing lignin–like compounds with O/C ratios of 0.1–0.4 and hydrogen/carbon (H/C) ratios of 0.7–1.5. (Yin et al., 2023). Despite varying removal efficiencies, these studies consistently demonstrate the A/O process’s ability to degrade and transform refractory organic matter, supporting its potential application in treating livestock and poultry digestate.

The integrated fixed–activated sludge (IFAS) process is another biological treatment technology that has shown great potential in degrading refractory organic matter. By combining suspended activated sludge with attached biofilm, IFAS promotes bacterial growth and enhances the degradation of nitrogenous compounds (Zhang et al., 2015). Ren et al. (2022) developed an IFAS process to mature landfill leachate and achieved a COD removal efficiency of 53% at an influent COD of 3,388 mg/L. At molecular level, Zhang et al. (2019) reported that IFAS process is particularly effective in degrading organic compounds with fewer carbonyl or carboxyl groups, higher aromatic content, and greater oxidative properties. In addition, organic compounds with greater saturation, lower molecular weight, and lignin–like structures are more easily adsorbed by extracellular polymeric substances (EPS) secreted by the biofilm, further enhancement of the removal effect of refractory organic matter (Zhang et al., 2019). These findings demonstrate the IFAS process’s effectiveness in removing and transforming refractory organics—especially aromatic compounds—highlighting its potential for treating livestock and poultry digestate (Zhang et al., 2023; Zhou et al., 2024).

Recent research has explored the integration of multiple biological treatment technologies to enhance the removal and transformation of refractory organic matter. The A/O process has been coupled with microbial fuel cells, membrane bioreactors, and sequencing membrane batch reactors to improve the degradation of refractory organic matter (Elmaadawy et al., 2020; Han et al., 2018; Liu et al., 2017; Zhang et al., 2019). For example, a study integrated the lab–scale A/O and Anammox processes to treat coal liquefaction wastewater (Zhang et al., 2019). Similarly, the IFAS system has been combined with sequencing batch reactors, membrane bioreactors, and moving bed biofilm reactors to enhance pollutant removal efficiency (Zhou and Xu, 2019; Bilad et al., 2011; Jin et al., 2013). Despite these advancements, limited studies have investigated the coupling of A/O and IFAS processes. Zhou et al., (2024a) developed an A/O–IFAS system and reported that the molecular weight, complexity, biodegradability, and degree of unsaturation of organic matter exhibited opposite trends following the A/O and IFAS treatment. Based on these previous studies, it is reasonable to hypothesize that the A/O and IFAS processes are complementary and can synergistically improve the removal and transformation of refractory organic matter during the treatment of livestock and poultry digestate.

In this study, the A/O and IFAS processes were coupled to treat real livestock and poultry digestate with the following objectives: (1) compare the efficiency of the two processes in removing refractory organic matter from livestock and poultry digestate; (2) characterize the transformation of refractory organic matter by the two processes; and (3) understand the mechanisms for the removal and transformation of refractory organic matter in the A/O and IFAS processes. To achieve these objectives, pilot–scale A/O and IFAS reactors were constructed, and the removal performance was evaluated with different operational conditions. Fluorescence spectroscopy, mass spectrometry, and microbial community analysis were conducted to further understand the transformation of refractory organic matter. The findings of this study are expected to provide valuable insights into the degradation mechanisms of refractory organic matter and the development of effective technologies for treating livestock and poultry digestate.

## 2. Materials and Methods

### 2.1 Reactor set–up and operation

The coupled system operated in continuous mode and primarily consisted of an A/O reactor and an IFAS reactor. The A/O reactor (50 L) was divided into four equal chambers. The first two were maintained under anaerobic conditions, while the latter two were aerated to sustain a dissolved oxygen (DO) concentration of 1 mg/L. The IFAS reactor (110 L) was similarly divided, with each chamber maintaining DO below 0.2 mg/L and equipped with mature biofilm carriers (32% filling ratio). A schematic diagram of the system is provided in Figure S1 of the Supporting Information (SI).

Seed sludge for the A/O reactor was conventional activated sludge (mixed liquor suspended solids, MLSS ∼3,500 mg/L), while the IFAS reactor was inoculated with sludge obtained from a system treating high–ammonia wastewater (MLSS ∼1,800 mg/L). All sludge and carriers were sourced from the Gaobeidian and Xiaohongmen wastewater treatment plants in Beijing, China.

Livestock and poultry digestate used as influent was collected from a farm in Rushan, Shandong Province, China. Its characteristics were as follows: COD, 2,258–6,850 mg/L; BOD₅, 845–1,062 mg/L; TN, 583–2,940 mg/L; alkalinity, ∼13,000 mg CaCO₃/L; and pH, 8.2–8.6. The A/O reactor received the raw digestate, whereas the IFAS reactor was fed with a diluted stream. Operational conditions for the IFAS reactor are summarized in Table 1, with additional details provided in SI Text S1.

**Table 1.** Operational conditions of the IFAS reactor.

| Phase | Unit | Phase I | Phase II | Phase III |
| --- | --- | --- | --- | --- |
| Time | d | 25–104 | 105–182 | 183–228 |
| Influent | – | 10%–30%<br>diluted A/O<br>reactor effluent | 40%–90% diluted<br>A/O reactor<br>effluent | 100% A/O reactor<br>effluent (undiluted) |
| Recirculation ratio | % | 400–550 | 300–700 | 200 |
| HRT | d | 3.0 – 4.0 | 4.0 – 4.5 | 4.0 – 1.0 |
| COD<sub>inf</sub> | mg/L | 270 – 1500 | 732 – 2840 | 1896 – 2258 |
| OLR | kg COD/(m<sup>3</sup>·d) | 0.1 – 0.4 | 0.2 – 0.7 | 0.5 – 0.7 |
| Average COD/N | – | 0.7 | 1.6 | 3.0 |

HRT: hydraulic retention time; OLR: organic matter loading rate

### 2.2 Sample collection and biochemical analysis

Water samples were collected from multiple points of the coupled system, including the influent, the second and fourth chambers of the A/O reactor, the primary settling tank, the regulating tank, the second and fourth chambers of the IFAS reactor, and the secondary settling tank (SI Figure S1). One sample was collected from each sampling point. COD, BOD_5_, TN, and alkalinity were measured following standard methods (APHA, 2005). Temperature and pH were determined using a water quality analyzer (WTW Multi 3420, WTW, Germany).

For EPS analysis, 25 mL of floc sludge samples were collected from the anaerobic and oxic end chambers of the A/O reactor and mixed as one representative sample. Similarly, 25 mL of floc sludge was collected from the second and fourth chambers of the IFAS reactor and mixed. Biofilm samples were obtained from the same positions, where the biofilm attached to the carrier was washed off with deionized water and pooled to a final volume of 50 mL. After sample collection, EPS was extracted using a modified heat extraction method (Wang et al., 2020b). The concentration of polysaccharides (PS) was measured using the anthrone method, with glucose as a standard (Frølund et al., 1996). The contents of proteins (PN) were analyzed using the Lowry method (Bradford, 1976).

### 2.3 Spectroscopic analysis

To identify fluorescent compounds in the water samples, excitation–emission matrix (EEM) fluorescence spectroscopy was performed using a multiwavelength fluorescence spectrophotometer (F7000, Hitachi, Japan). Milli–Q deionized water was used as a blank control for each sample and subtracted from the measurements. The excitation and emission wavelengths were scanned over the range of 220–550 nm with a 5 nm interval.

EEM data were analyzed using parallel factor analysis (PARAFAC) in MATLAB (version 2016a) with the DOMFluor toolbox (Stedmon and Bro, 2008). A PARAFAC model (n = 60) was applied to extract fluorescence components, and their relative proportions were determined using the maximum fluorescence intensity (F_max_). These components were then compared with literature data via OpenFluor, based on excitation (Ex) and emission (Em) wavelength similarities. Model validation was performed using split–half analysis and residual analysis.

To further explore trends in identified fluorescent compounds, two–dimensional correlation spectroscopy (2D–COS) was applied to the F_max_ data obtained from EEM– PARAFAC (Yu, 2019). All computations were performed in MATLAB, following the method described by Noda and Ozaki (Noda and Ozaki, 2005). Additional details on the 2D–COS are provided in SI Text S2.

### 2.4 Mass spectrometric analysis

Fourier transform–ion cyclotron resonance mass spectrometry (FT–ICR MS) was employed to characterize the molecular composition of dissolved organic matter (DOM) (Z. Chen et al., 2024). DOM was extracted and concentrated using a solid–phase extraction method, as detailed in SI Text S3. The dried extracts were analyzed using a Bruker 15T FT–ICR mass spectrometer (Germany) equipped with an electrospray ionization source operating in negative ion mode.

The molecular composition of refractory organic matter was visualized using a Van Krevelen diagram, in which the H/C ratio for each formula was plotted versus its O/C ratio. The Van Krevelen diagram of this study was divided into seven subclasses (Hockaday et al., 2009): lipids (H/C =1.5–2.0, O/C = 0–0.3), carbohydrates (H/C =1.5–, O/C = 0.67–1.2), proteins (H/C = 1.5–2.2, O/C = 0.3–0.67), lignins (H/C = 0.7–1.5, O/C = 0.1–0.67), aromatic structures (H/C = 0.2–0.7, O/C = 0–0.67), unsaturated hydrocarbons (H/C = 0.7–1.5, O/C = 0–0.1), and tannins (H/C = 0.6–1.5, O/ C = 0.67– 1.0).

A series of molecular indices were calculated to further characterize variations in organic matter composition (Wang et al., 2021). These included molecular relative intensity (M_i_), weight–averaged molecular weight (MW_wa_), double bond equivalents (DBE), average DBE (DBE_wa_), DBE minus oxygen per carbon ((DBE−O)/C), nominal oxidation state of carbon (NOSC), average NOSC (NOSC_wa_), average atomic ratios of oxygen to carbon (O/C_wa_) and hydrogen to carbon (H/C_wa_), modified aromaticity index (AImod), and molecular lability index (MLB_L_). The corresponding equations are provided in Supporting Information Text S4.

To gain in–depth insight into the transformation of DOM, molecular detected by FT– ICR MS were classified into three distinct categories: precursors, resistants, and products. The approach is detailed in SI Text S5. Reaction pathways were further elucidated by constructing mass difference networks using an established R script (Lin et al., 2024). The results were visualized using Gephi software (version 0.10), a freely accessible tool for network analysis.

### 2.4 Microbial community analysis

Biomass samples were collected from the anaerobic and oxic end chambers of the A/O reactor and the second and fourth chambers of the IFAS reactor (SI Figure S1). DNA was extracted using a FastDNA Spin Kit for Soil (QBIOgene Inc., Carlsbad, CA). Sequencing of the 16S rRNA gene was performed by Majorbio Technology Co., Ltd. (Shanghai, China) with an Illumina MiSeq platform. The bacterial 16S rRNA gene primer pair was 338F (5 ′ – ACTCCTACGGGAGGCAGCAG – 3 ′) and 806R (5 ′ – GGACTACHVGGGTWTCTAAT–3′) for the V3 ∼ V4 region. Paired–end sequences were assembled and denoised using QIIME 2, and operational taxonomic units (OTUs) were picked using DADA2 (Callahan et al., 2016; Caporaso et al., 2010). Taxonomy was assigned using the QIIME 2 plugin feature–classifier and the SILVA database as the reference, with the classifier trained on 99% OTUs (Quast et al., 2013).

Phylogenetic Investigation of Communities by Reconstruction of Unobserved States (PICRUSt2) was used to predict the overall genetic potential of the microbial communities. The prediction was made by using 16S rRNA sequencing and referenced Kyoto Encyclopedia of Genes and Genomes (KEGG) database. To compare the differences of metabolic pathways in A/O and IFAS processes, Statistical Analysis of Metagenomic Profiles (STAMP, version 2.1.3) was used to analyze KEGG prediction results based on 16S rRNA sequences.

### 2.5 Statistical analysis

All statistical analyses were performed using the software R. Prior to hypothesis testing, data were assessed for normality using the Shapiro–Wilk test. A one sample t–test was conducted to determine whether the sample mean significantly differed from a specified reference value. If the normality assumption was violated, the Wilcoxon signed–rank test was used as a non–parametric alternative. A *p*–value < 0.05 was considered statistically significant (Yao et al., 2024).

To explore the relationship between microbial community composition and DOM characteristics, Mantel tests were conducted based on Bray–Curtis and Euclidean distance matrices (Wang et al., 2021). Microbial beta diversity was calculated from 16S rRNA gene–based OTU abundance profiles using Bray–Curtis dissimilarity. Euclidean distance matrices for DOM were constructed using molecular indices derived from FT– ICR MS and F_max_ from EEM–PARAFAC. Mantel tests were performed using the “vegan” package in R with 9,999 permutations, and correlations with *p*–values < 0.05 were considered significant.

## 3. Results and Discussion

### 3.1 Organic removal

Real livestock and poultry digestate entering the A/O process had a high COD up to 6,850 mg/L (COD_A/O–inf_, Figure 1A) but a low BOD_5_ (<1,062 mg/L, Table 1), highlighting the refractory nature of the organic matter. As the A/O process can typically withstand fluctuating organic loading (Cheng et al., 2020), the A/O reactor was fed with undiluted digestate, and its COD removal performance was optimized by varying the HRT and aeration rate. The functional microorganisms in IFAS was more sensitive to organic loading shock (Li et al., 2024). Therefore, the IFAS reactor was fed with diluted A/O effluent, and its COD removal performance was optimized by varying the HRT and dilution ratio (Table 1).

**Figure 1.**
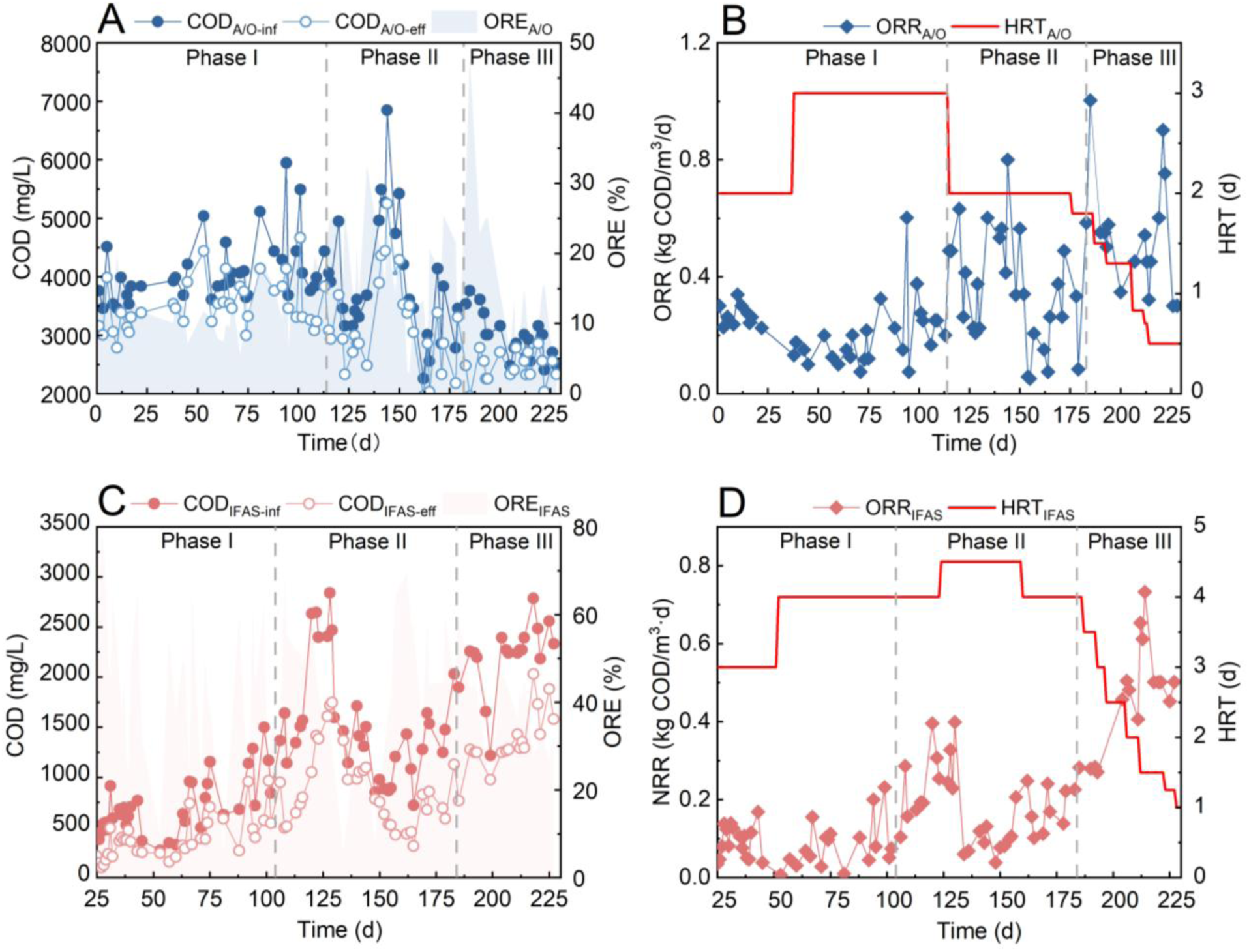
COD removal performance of the A/O and IFAS reactors. (A) COD_inf_, COD_eff_ and ORE of the A/O reactor; (B) ORR and HRT of the A/O reactor; (C) COD_inf_, COD_eff_ and ORE of the IFAS reactor; and (D) ORR and HRT of the IFAS reactor.

The HRT of the A/O reactor increased from 2 to 3 days in Phase I (Days 0–114), while the aeration rate was maintained at 20 L/h. As a result, the average organic removal efficiency (ORE) decreased slightly from 14% with 2–d HRT to 13% with 3–d HRT (t– test, *p* < 0.05, Figure 1A). Similarly, the average organic removal rate (ORR) decreased from 0.3 kg COD/(m³·d) with 2–d HRT to 0.2 kg COD/(m³·d) with 3–d HRT (t–test, *p* < 0.05, Figure 1B). Because the HRT of 2 d showed a favorable efficiency and rate, it was selected to understand the effect of aeration rate on removal performance in Phase II (Days 115–182). The aeration rate was maintained at 40 L/h for 30 d and then increased to 60 L/h. As a result, the average ORE rose from 15% to 20% (t–test, *p* < 0.05), and the average ORR was significantly improved from 0.2 kg COD/(m³·d) to 0.4 kg COD/(m³·d) (t–test, *p* < 0.05). These results, which are consistent with previous studies (Luanmanee et al., 2002; Wang and Zhang, 2017), suggest that higher oxygen supply positively affects the performance of the A/O process.

For the IFAS reactor, within the first 30 days of Phase I (Days 25–104), the influent was 10% A/O effluent and 90% water to achieve rapid startup. At this stage, the average ORE was 49%, while the ORR remained low at 0.1 kg COD/(m³·d). After 55 days, the proportion of A/O effluent was progressively increased to 30%, and the average influent COD concentration rose to 841 mg/L. Meanwhile, the HRT was extended from 3 days to 4 days to alleviate the potential inhibitory effects of refractory organic matter on IFAS. By the end of this phase, the ORE decreased to 41%, while the ORR remained stable at 0.1 kg COD/(m³·d). In Phase II (Days 105–182), the proportion of A/O effluent in the IFAS influent was further increased from 40% to 90% by 10% increments, resulting in an final influent COD of 1,521 mg/L, an average ORE of 41%, and an average ORR of 0.2 kg COD/(m³·d). Notably, between Days 120 and 129, an abrupt rise in influent COD concentration to 2,566 mg/L led to a temporary decline in ORE. In response to the presence of high contractions of refractory organic matter, the HRT was extended to 4.5 days for seven days and reduced back to 4 days, after which the IFAS reactor achieved an ORE of 52% while maintaining an ORR of 0.2 kg COD/(m³·d).

Because the IFAS reactor could function under high and fluctuating organic loading, Phase III (Days 183–228) was started by feeding the effluent from the A/O reactor directly into the IFAS reactor without dilution. Meanwhile, the HRT of the A/O and IFAS reactors gradually reduced to 0.5 d and 1 d, respectively, to accelerate organic removal (Figures 1B and 1D). Consequently, the average ORE of the A/O and IFAS reactors reached 17% and 43%, respectively. Both reactors achieved an average ORR of 0.5 kg COD/(m³·d), significantly higher than that in the first two phases. The results suggest that the A/O and IFAS reactors are successfully optimized to treat real livestock and poultry digestate with satisfactory efficiencies and rates. Compared to previous studies conducted with small–scale reactors (2 – 22 L), low organic loading (<2,000 mg/L), and high BOD_5_/COD ratio (Table 2), this study achieved a larger effective volume and relatively faster removal of real livestock and poultry digestate. With an overall ORE of 60% after A/O and IFAS, the coupled system demonstrated effective degradation of refractory organic matter, highlighting its potential for practical applications in treating high–strength low–biodegradability wastewater.

**Table 2.**
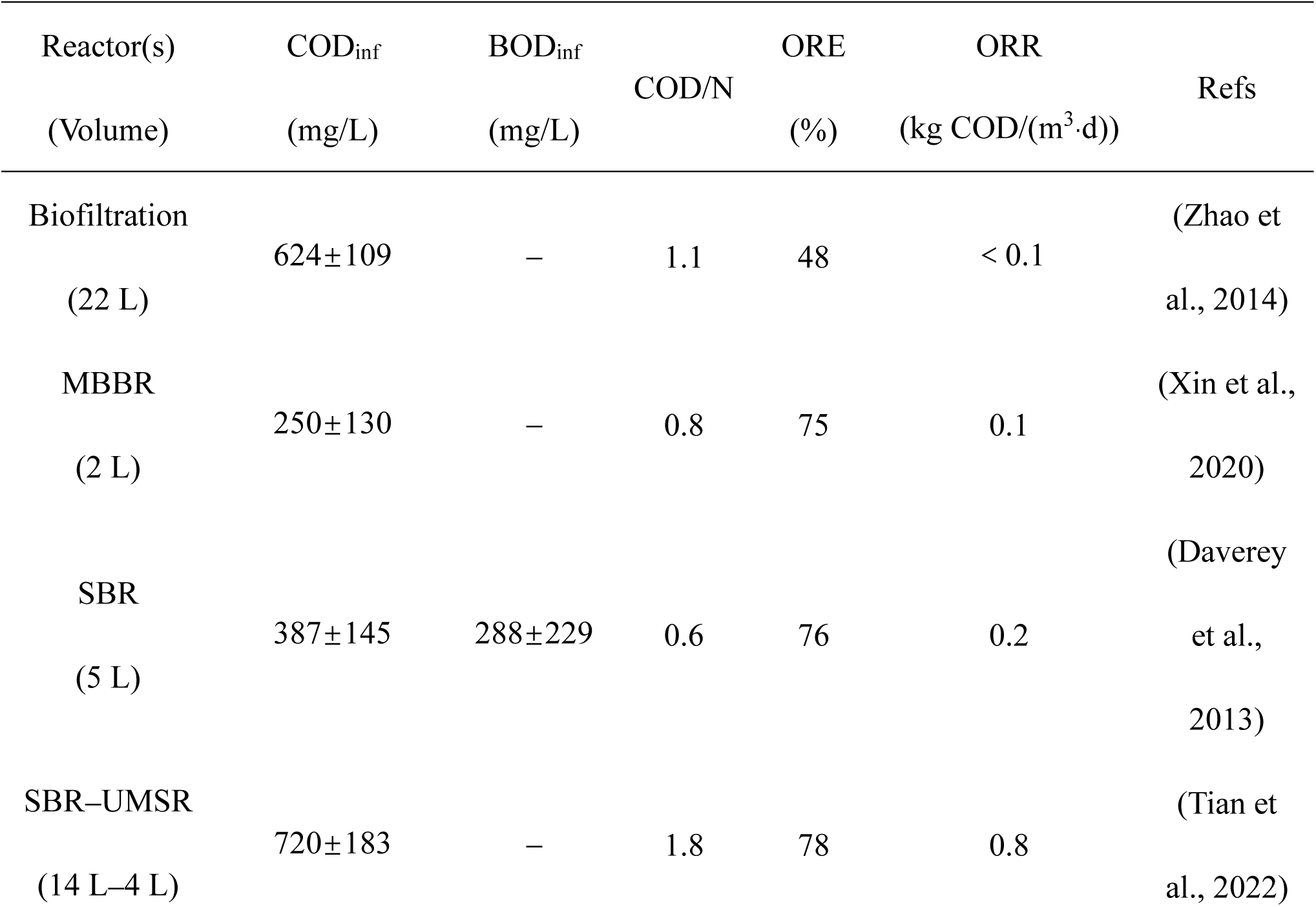
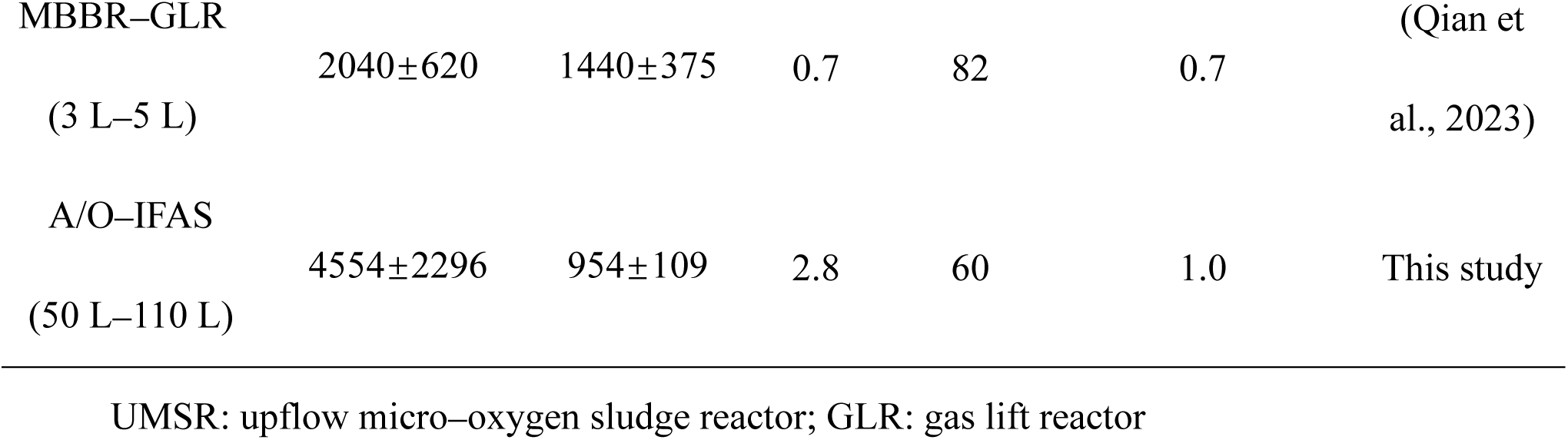
Performance of livestock and poultry digestate treatment using biological process.

### 3.2 Organic transformation revealed by fluorescence spectroscopy

To determine the composition and fate of the organic matter in livestock and poultry digestate, samples at different treatment stages were collected and analyzed using EEM fluorescence spectroscopy (SI Figure S2). Based on the PARAFAC model, three distinct humic–like refractory components were identified (Figure 2). Component 1 (E_x_/E_m_ = 410/490 nm), which was previously reported by Stedmon and Markager, 2005 (Figure 2A), might originate from terrestrial sources and undergo more significant degradation during biological treatment than the other components. Component 2 (E_x_/E_m_ = 365/440 nm), which was aligned with the one reported by Chen et al., 2018, could also be attributed to terrestrial origin and was commonly associated with recycled water and wastewater discharge (Figure 2B). Component 3 (E_x_/E_m_ = 285, 310/410 nm) resembled the microbial–derived organic matter reported by Hong et al., 2021 (Figure 2C).

**Figure 2.**
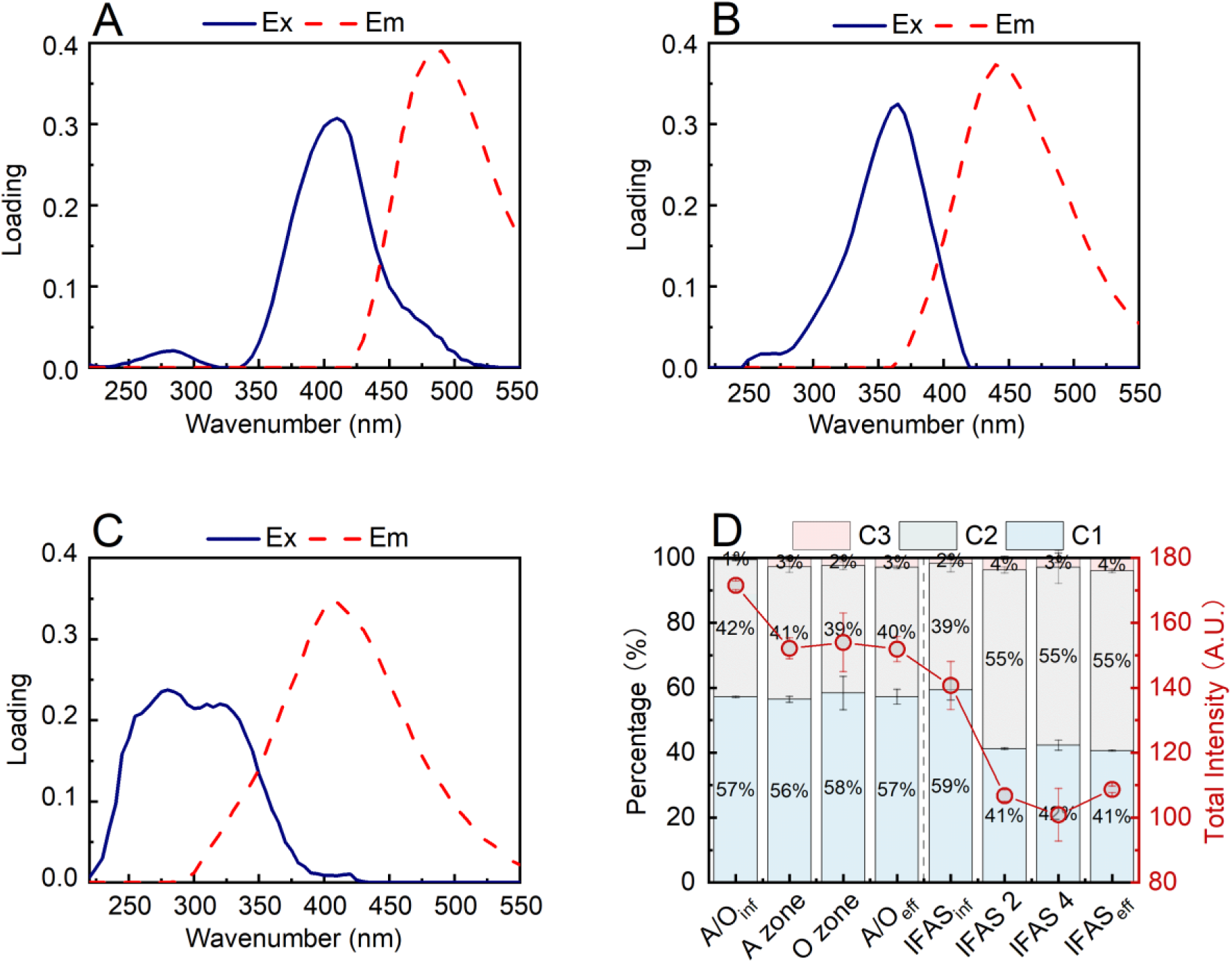
(A) Component 1, (B) Component 1, and (C) Component 3 identified using the PARAFAC model. (D) Variation of fluorescent components in the A/O and IFAS processes.

Although all three components were classified as humic–like substances, their distinct characteristics resulted in different degradation patterns in the A/O and IFAS processes (Figure 2D). The A/O effluent showed minimal compositional ratio changes compared to its influent, suggesting non–selective removal of the three components (Zhang et al., 2019). In contrast, the IFAS process significantly altered the effluent composition by selectively removing Component 1. A more detailed inspection revealed that Component 1 was the dominant fraction in the influent for both processes, accounting for up to 57% and 59%, respectively (Figure 2D). The A/O unit removed 11% of this component as evidenced by the change in the overall fluorescence intensity, whereas the IFAS unit achieved substantially higher removal efficiency of 47%. Component 2 followed the opposite trend, with 16% removed in the A/O process but a 10% increase in the IFAS effluent, likely due to its generation during treatment. Component 3 showed the lowest abundance in raw digestate (<1%) but increased significantly after the treatment, with its concentration increased by 4 times in the A/O effluent and 1.8 times in the IFAS effluent.

2D–COS analysis was combined with PARAFAC modeling to determine the sequential variations within each component (Yu, 2019) (Figure 3). The auto–peaks on the diagonals of the synchronous maps revealed that Component 1 in the digestate was the most susceptible to degradation in both the A/O and IFAS processes (Figures 3A and 3D), consistent with the findings by Stedmon and Markager, 2005. On the other hand, Component 3 was the least responsive to the treatment (Figure 3D) (Noda and Ozaki, 2005), possibly due to its initially low content in the real digestate. The asynchronous maps (Figures 3B and 3E) and the resulting cross–peak signs (Figures 3C and 3F) indicated a consistent reaction priority in both the A/O and IFAS reactors following the order of Component 2, Component 3, and Component 1 (Noda and Ozaki, 2005). Despite this similarity, as revealed by Figure 2D, the A/O process was slightly effective at removing Component 2 than other components, while the IFAS process demonstrated effective degradation of Component 1 but led to the accumulation of Component 2. These findings highlight the distinct roles of the A/O and IFAS processes in transforming the refractory organic matter in livestock and poultry digestate.

**Figure 3.**
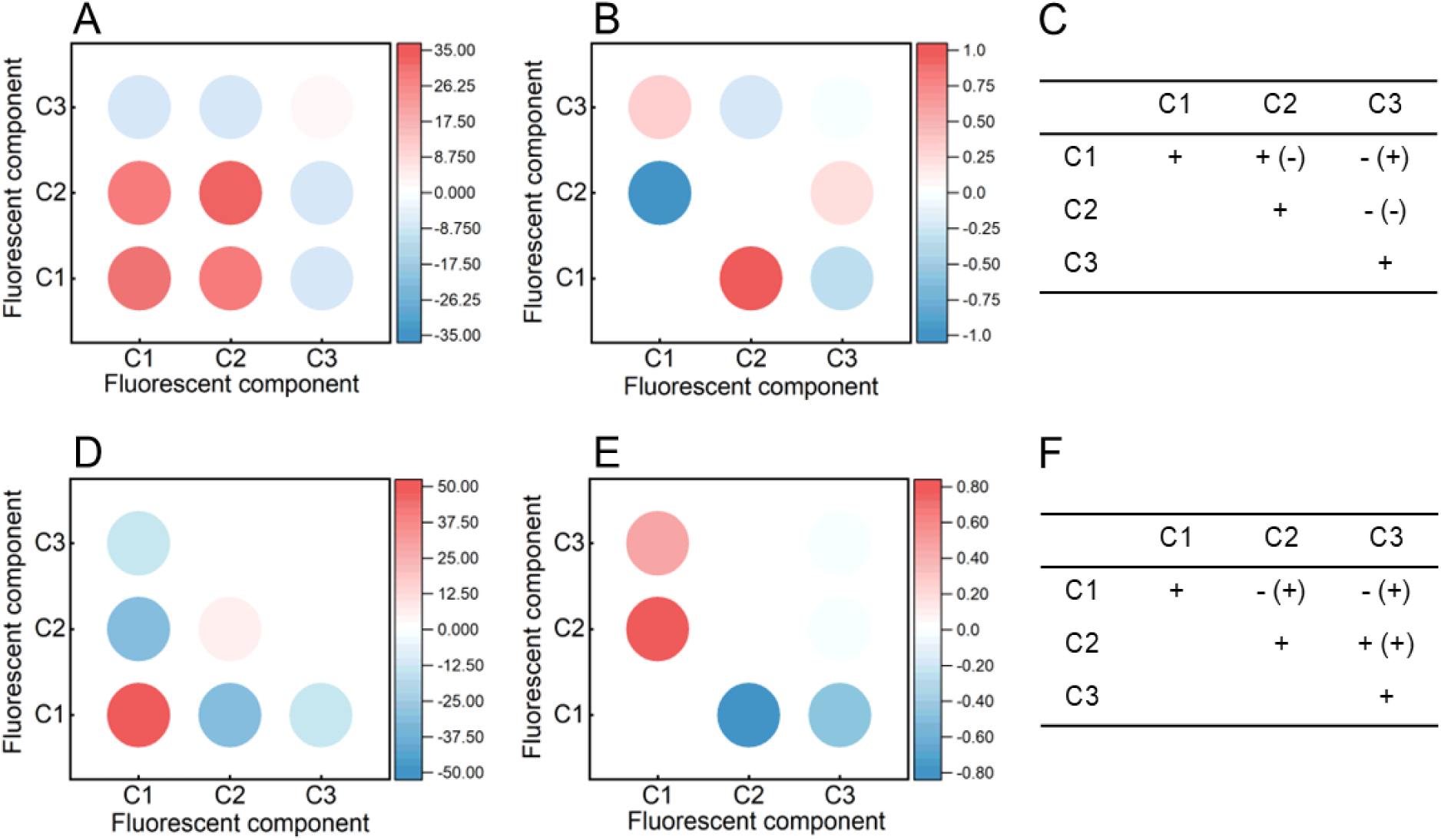
(A) Synchronous maps, (B) Asynchronous maps, and (C) Signs generated from EEM– PARAFAC–COS of the A/O process. (D) Synchronous maps, (E) Asynchronous maps, and (F) Signs generated from EEM–PARAFAC–COS of the IFAS process. Signs were obtained from the upper–left corner of the maps.

### 3.3 Organic transformation revealed by mass spectrometry

#### 3.3.1 Variation in molecular characteristics

Given the complexity of biological treatment of real livestock and poultry digestate, molecular–level analysis was conducted to further understand the transformation of organic matter. The changes in organic molecular composition throughout the treatment processes, along with the calculated molecular indices, are presented in SI Figure S3 and Table S1, respectively. Distinct trends in both the number of organic formulas and their molecular weight were observed following the A/O and IFAS treatments. Although molecules with molecular weights ranging from 300 to 500 Da dominated across both processes (SI Figure S4A), the A/O process resulted in an increase in both the number of detected formulas and average molecular weight. In contrast, the IFAS process led to a reduction in these parameters (SI Table S1). These findings differ from previous research on municipal wastewater (Zhou et al., 2024), likely due to differences in influent composition and the recalcitrant nature of livestock–derived organic matter.

To further elucidate the transformation of organic matter, molecular indicators reflecting chemical properties, including DBE, NOSC, and AI_mod_, were analyzed. DBE signifies the number of double bonds or rings in a molecule and can reflect molecular reactivity. During the A/O process, DBE remained largely unchanged, suggesting stable molecular reactivity throughout the treatment. In contrast, the IFAS treatment caused an increase in the proportion of higher–DBE molecules (SI Figure S4B), indicating more active transformation including addition, elimination, and/or redox reactions (Lin et al., 2024). NOSC can serve as a thermodynamic indicator of the biodegradability of organic molecules (LaRowe and Van Cappellen, 2011). In both A/O and IFAS treatments, the number of molecules with –1 < NOSC < 0 (typically hydrocarbons) decreased, while the number of molecules with NOSC > 0 (typically carbon oxides) increased (Lin et al., 2024; Zhou et al., 2019) (SI Figure S4C), implying the degradation of hydrocarbons to carbon oxides. The AI_mod_ index reflects the density of unsaturated carbon bonds and molecular aromaticity (Koch and Dittmar, 2006). Following the treatment, the AI_mod_ values remained stable after A/O but increased after IFAS (SI Figure S4D). The higher level of molecular complexity and aromaticity after IFAS aligned with a previous study (Zhang et al., 2019).

#### 3.3.2 Possible precursor–product pairs

Van Krevelen diagram analysis was conducted to classify molecules into precursors, products, and resistant compounds (Smith et al., 1994). As shown in Figures 4A and 4B, lignin–like molecules exhibited the most substantial changes, accounting for over 50% of all molecular alterations in both the A/O and IFAS processes. Protein–like molecules accounted for the second–largest category, comprising 15% of the transformed molecules in the A/O process and 18% in the IFAS process (SI Figure S5). Overall, the A/O process contained less precursor molecules but more products and resistant compounds compared to the IFAS process (SI Figure S5), consistent with a previous study by Zhang et al. (2019).

**Figure 4.**
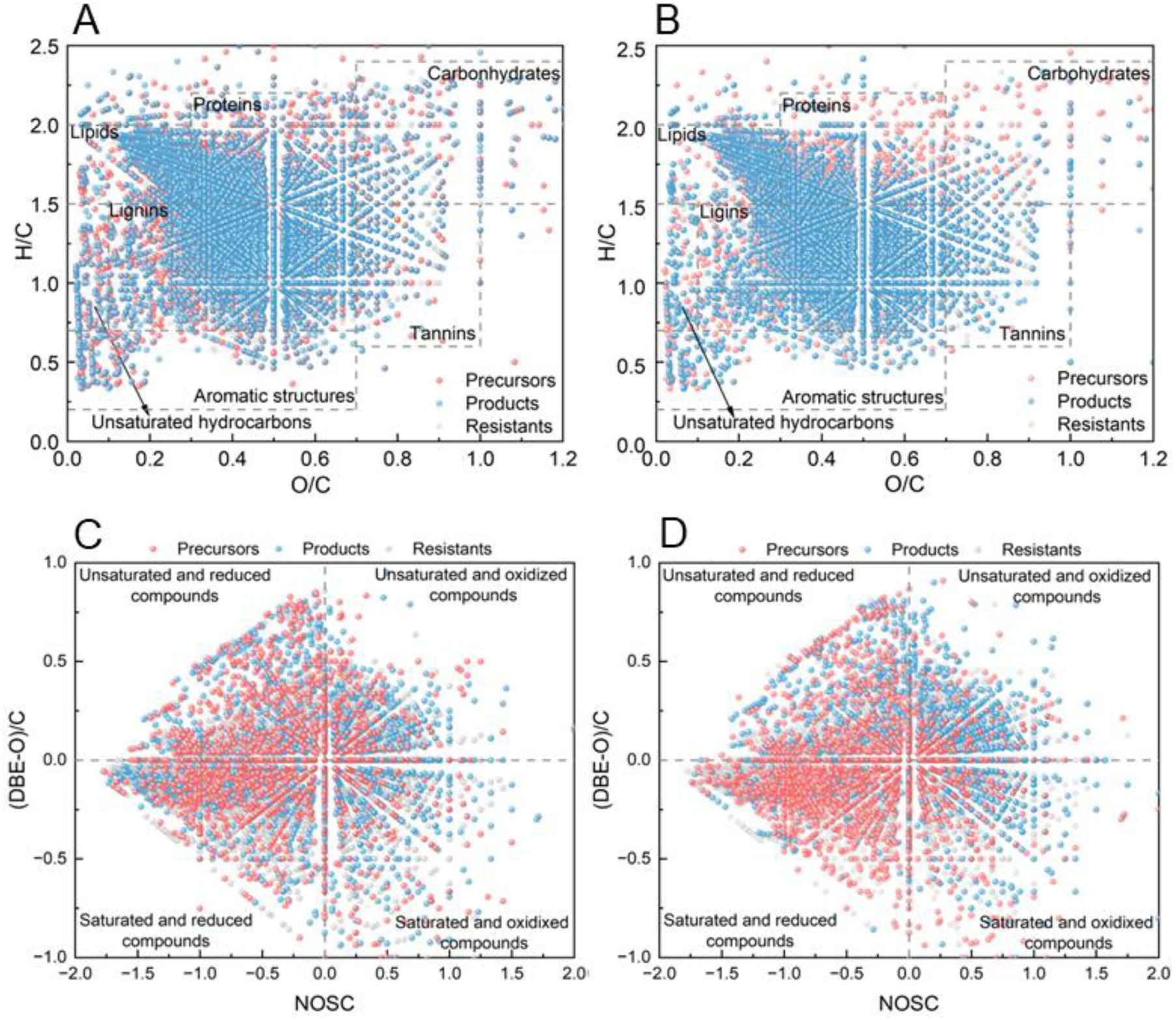
Van Krevelen diagrams classifying molecules into precursors (red dots), products (blue dots) and resistants (grey dots) DOM. Classification based on organic matter species in the (A) A/O process and (B) IFAS process. Classification based on chemical properties (C) A/O process and (D) IFAS process.

(DBE−O)/C and NOSC values were used to further categorize individual molecules into four groups: unsaturated and oxidized, unsaturated and reduced, saturated and reduced, and saturated and oxidized (Figures 4C and 4D). The A/O processes showed superior performance in altering molecules in the reduced state, with up to 65% of all precursors being transformed. However, the A/O process showed no significant difference in the removal of saturated and unsaturated molecules and achieved a 50 % conversion for both categories. In contrast, the IFAS process exhibited superior conversion performance for molecules in the saturated and reduced states, accounting for 42% of all precursors. In terms of products, the A/O process generated more reduced substances, accounting for 57% of all products, while the IFAS process predominantly produced molecules in the unsaturated state, accounting for 65%. Overall, the A/O process excelled in converting and producing reduced molecules, while the IFAS process promoted the transformation of saturated and reduced precursors and the generation of unsaturated products.

Mass difference network analysis was conducted to correlate the precursor–product pairs across 32 identified reactions (Schollée et al., 2018; Zhang et al., 2021). Among all molecular groups, CHONs were the most frequently transformed compounds in both the A/O and IFAS processes, contributing to 31% and 28% of the total transformations, respectively (Figures 5A and 5B). CHONSs were the second most abundant molecular group involved in transformations, accounting for 24% and 25% in the A/O and IFAS processes, respectively. These findings highlight the crucial role of nitrogen and sulfur in the transformation pathways of refractory organic matter. From the perspective of reaction types, oxygenation was the predominant pathway in the A/O process (∼29% of the total reactions) followed by dealkylation (28%). In contrast, the IFAS process exhibited a higher prevalence of dealkylation (31%) and oxygenation (22%). Among the identified precursor–product pairs, demethylation (–CH_2_) was the most frequently observed transformation in the A/O process, while dehydrogenation (–2H) was the dominated transformation in the IFAS process, with both reactions accounting for 7% of the total transformations.

**Figure 5.**
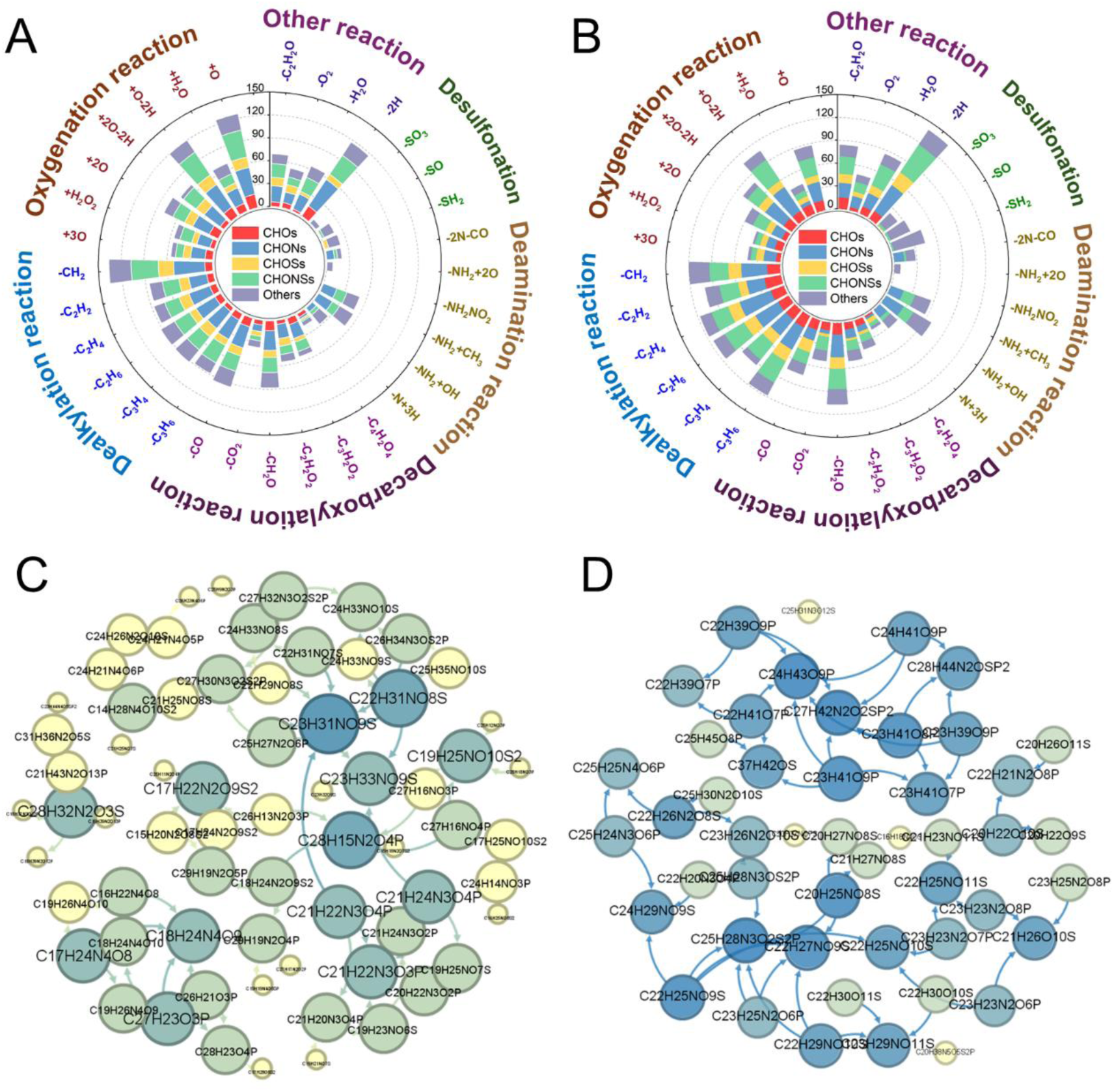
Radar plots showing the number of possible precursors–product pairs during the (A) A/O and (B) IFAS processes. The scales in the radar plots indicate the total number of reactions observed, while labels represent specific chemical transformations occurred. Molecular network analysis of the organic molecules in the (C) A/O and (D) IFAS process. The arrows in the networks denote the direction of molecular transformations, and darker colors and larger nodes indicate molecules involved in a greater number of transformation pathways.

The molecular network revealed a total of 2,739 and 2,685 transformations of organic molecules were identified in the A/O and IFAS processes, respectively (Figures 5C and 5D). Approximately 55% of the precursor molecules in the A/O process and 53% in the IFAS process participated in the formation of multiple products. Notably, 11% of precursors in the A/O process and 8% in the IFAS process were involved in more than three distinct transformation pathways, each leading to the generation of different product These results highlight the high degree of molecular reactivity and structural diversity, as well as the complexity of organic matter transformation during biological treatment. In the A/O process, the most predominant precursor C_28_H_15_N_2_O_4_P participated in the formation of 11 different products. Other versatile precursors, such as C_27_H_32_N_3_O_2_S_2_P, C_27_H_16_NO_3_P and C_27_H_16_NO_4_P, were involved in the generation of 10 products. Primary products in the A/O process, such as C_18_H_24_N_4_O_9_, C_23_H_3_1NO_9_S, and C_23_H_33_NO_9_S, originated from nine different precursors. In the IFAS process, C_22_H_25_NO_9_S was the predominant precursor, contributing to the formation of 10 products. This molecule had relatively lower carbon numbers and unsaturation compared to the precursors in the A/O process. Meanwhile, C_15_H_18_N_4_O_9_ and C_24_H_33_NO_10_S were the most frequently observed products in the IFAS process, each derived from more than 10 different precursors. These findings highlight the distinct roles of specific precursor molecules and their diverse transformation pathways in the A/O and IFAS processes.

### 3.4 Microbial community and activity

The removal and transformation of refractory organic matter are largely dependent on the composition of the microbial communities and the activity of functional populations (Wang et al., 2021). As shown in Figure 6A, the A/O communities were dominated by *Firmicutes* (relative abundance 36%) and *Proteobacteria* (28%), whereas the IFAS communities were dominated by *Chloroflexota* (35%) and *Proteobacteria* (23%). *Firmicutes* is a widely distributed and functionally diverse phylum, in which some members have been reported to degrade polysaccharides and complex organic compounds in anaerobic digestate (Fan et al., 2023; Lu et al., 2024). *Proteobacteria*, a highly diverse bacterial phylum, comprises many well–characterized members that can break down a wide range of organic pollutants, including hydrocarbons, phenols, aromatic compounds, and polycyclic aromatic hydrocarbons (Haritash and Kaushik, 2009; Zhang et al., 2024). A previous study also reported the enrichment of *Chloroflexota* under long–term exposure to wastewater with low biodegradability and their interactions with autotrophic microorganisms, such as *Planctomycetota*, to facilitate the degradation of refractory organic matter (Wen et al., 2021). Indeed, *Planctomycetota* was observed with a high abundance of 9% in IFAS process but not in the A/O process. The difference in microbial community composition in the two processes explained their different behaviors in organic removal and transformation (Figures 1–5).

**Figure 6.**
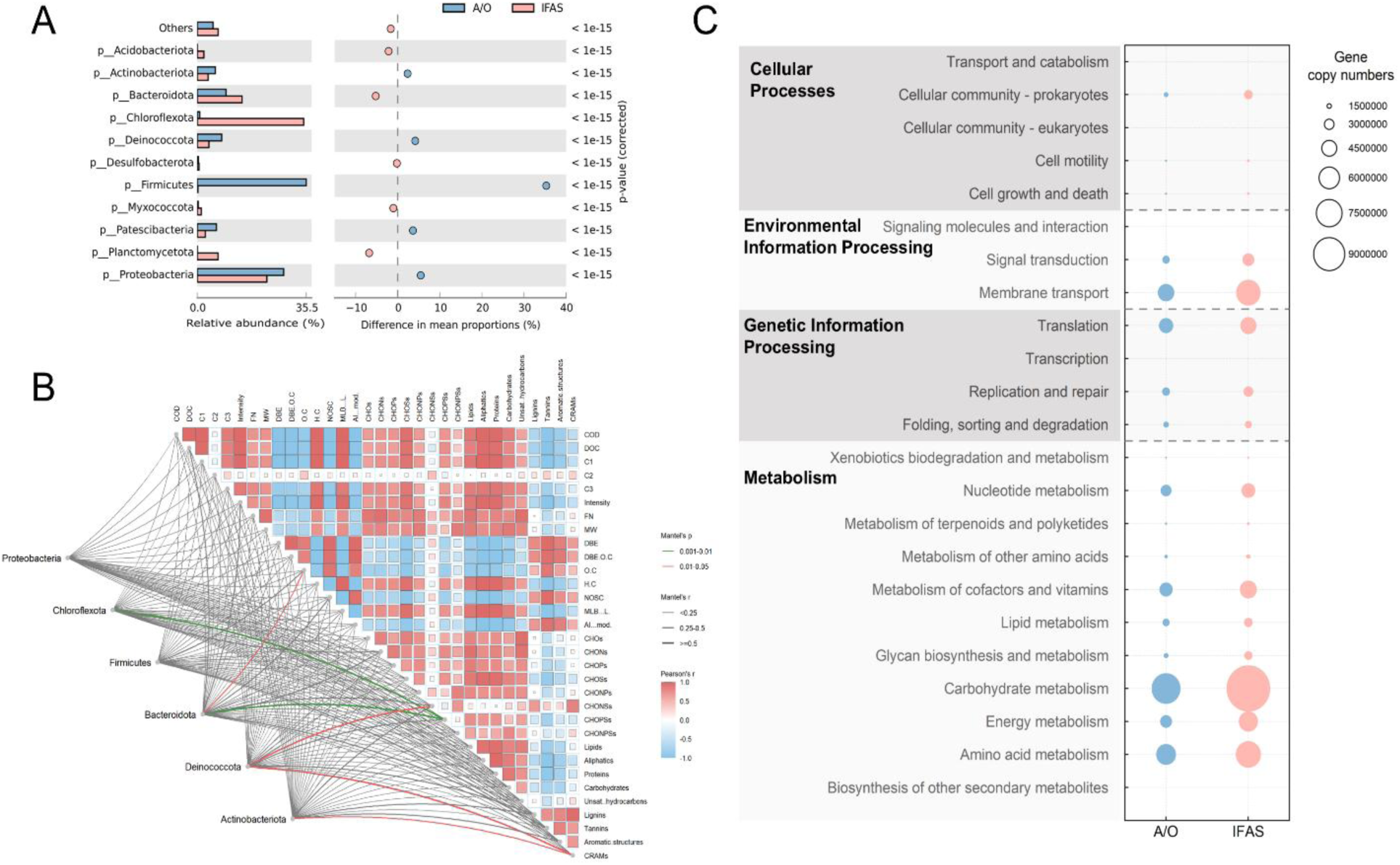
(A) Relative abundance of dominant phyla (left) and difference in relative abundance in the A/O and IFAS processes (right). (B) Correlation between dominant phyla, fluorescence spectroscopy results, and molecular variation. Solid lines represent the Mantel test results between phyla and organic matter characteristics, and the matrix depicts Pearson correlation coefficients between spectroscopic parameters and molecular variations. (C) Prediction of metabolic function at KEGG Level 2 based on microbial community composition.

Correlation analyses implied noticeable contribution of some of these phyla to organic removal and transformation (Figure 6B). For example, the abundances of *Chloroflexota*, *Bacteroidota*, and *Deinococcota* correlated with the concentrations of nitrogen–, sulfur–, and phosphorus–containing organic matter (*p* < 0.05) with Mantel’s r ranging from 0.26 to 0.80. These associations indicate potential involvement of these phyla in the transformation of heteroatom–containing organic matter. A strong positive correlation between nitrogen–sulfur and phosphorus–sulfur compounds further imply a shared origin and possible transformation through parallel microbial pathways, such as nitrogen/phosphorus substitution and sulfur oxidation. *Bacteroidota* was positively correlated with the O/C ratio of organic molecules (Mantel’s r 0.43), indicating a preference for oxygen–rich substrates such as carboxylated aromatics, hydroxylated aliphatics, and polysaccharide derivatives. These compounds typically require hydrolytic or oxidative enzymes for degradation. The presence of diverse carbohydrate–active enzymes and fermentative capacity in *Bacteroidota* supports its role in breaking down complex oxygenated organics, even under low–oxygen conditions (Martens et al., 2009). Additionally, a negative correlation was observed between the O/C ratio and both EEM–PARAFAC Components 1 and 3 as well as molecular weight, indicating that highly oxidized compounds tend to be smaller in size and exhibit weaker fluorescence signals. This trend is consistent with the oxidative breakdown of humic–like substances into low molecular weight, oxygen–rich degradation products (Lin and Guo, 2020). Together, these results highlight the role of *Bacteroidota* in converting high–molecular–weight, humic–like compounds into more oxidized, biodegradable molecules. *Deinococcota* and *Actinobacteria* showed strong correlations with carboxyl–rich alicyclic molecules (Mantel’s *r* = 0.68 and 0.38, respectively), which were positively associated with nitrogen– and sulfur–containing organics and lignin–like structures and negatively with COD concentration. This suggests that carboxyl–rich alicyclic compounds may act as intermediates formed during the microbial transformation of lignin–derived and heteroatom–containing organic matter. Known for their tolerance to oxidative stress and their ability to degrade aromatic compounds, *Deinococcota* may contribute to the cleavage and oxidation of chemically stable ring structures (Santos et al., 2019). A*ctinobacteria*, widely recognized for their ligninolytic enzyme systems, likely facilitate demethylation and ring–opening reactions in lignin–like compounds (Levy-Booth et al., 2022). The combined activity of these two phyla may promote the breakdown of structurally complex aromatic substances into smaller, more oxidized and biodegradable molecules, thereby enhancing COD reduction and driving the molecular transformation of DOM during the biological treatment of livestock and poultry digestate.

The overall activity of the communities was assessed by measuring the content of EPS, which could play a key role in the removal and transformation of refractory organic matter (Zhang et al., 2020). As illustrated in SI Figure S6, after acclimation, the EPS content significantly increased in both reactors, from 149 to 247 mg/g VSS in the A/O process and from 58 to 182 mg/g VSS in the IFAS process. This increase coincided with enhanced organic matter removal observed during the later operational stages (Figure 1), likely due to the promotion of adsorption–biodegradation processes mediated by EPS. Notably, the EPS produced in the A/O reactor exhibited a lower protein–to– polysaccharide (PN/PS) ratio (5.5) compared to that in the IFAS reactor (7.0). A lower PN/PS ratio indicates a smaller proportion of hydrophobic domains within the EPS matrix, resulting in fewer available binding sites for hydrophobic organic compounds (Chen et al., 2024). This may partially explain the relatively lower removal efficiency observed in the A/O process. Previous studies suggested that refractory organic matter stimulated the secretion of EPS (Miao et al., 2018). Negatively charged functional groups in EPS can electrostatically interact with positively charged refractory organic matter, while hydrophobic regions serve as adsorption sites for non–polar molecules (Sheng et al., 2010). Once adsorbed, these organics become more accessible to microbial enzymatic degradation (Liang and Liu, 2008). Therefore, EPS not only serve as a protective biofilm matrix but also function as an active interface that facilitates the capture and subsequent biodegradation of refractory organic matter during biological treatment.

The overall activity of the communities was further predicted using PICRUSt2 (Figure 6C) (Z. Chen et al., 2024). Carbohydrate metabolism and amino acid metabolism were inferred to be the predominant metabolic pathways in both processes. Notably, genes associated with these metabolic functions was inferred to have a higher copy number in the IFAS process, potentially explaining its superior performance in organic removal (Figure 1). Pathways related to carbohydrate metabolism and amino acid metabolism were highly enriched in the IFAS reactor. This suggests a stronger capacity for degrading labile organic matter, consistent with the higher abundance of *Bacteroidota*, a phylum well–known for degrading hydrolyze polysaccharides and other oxygen–rich substrates (Valdezate et al., 2015). The elevated O/C ratios observed in IFAS also support the preferential degradation of oxygenated compounds by *Bacteroidota*. The elevated functional genes related to membrane transport and signal transduction in the IFAS system also suggest enhanced microbial communication, substrate sensing, and uptake—critical for efficient adaptation to high–strength organic loading (Özkan et al., 2024). In parallel, enrichment in transcription, translation, and protein folding pathways indicates robust gene expression and protein biosynthesis, likely supporting the active degradation of both labile and recalcitrant DOM (Njenga et al., 2023). Collectively, these results highlight that the superior performance of the IFAS system is underpinned by both taxonomic enrichment and functional enhancement. The synergistic interaction between *Bacteroidota*, *Actinobacteria*, and *Deinococcota*, along with their associated metabolic capacities, enables the effective transformation of structurally diverse organic compounds—from readily biodegradable substrates to lignin–like recalcitrant fractions—thereby contributing to comprehensive organic matter removal in livestock and poultry wastewater treatment.

## 4. Conclusion

This study investigated the removal performance, molecular characteristics of organic matter, and microbial community structures in the A/O and IFAS processes during the treatment of livestock and poultry digestate. The key conclusions drawn from this research are summarized as follows:

1. Both the A/O and IFAS processes effectively utilize refractory organic matter from livestock and poultry digestate as carbon sources. However, the IFAS process demonstrated a higher removal efficiency, especially in degrading molecules with highly aromatic structures.
2. The A/O process showed greater efficiency in converting and producing reduced molecules, predominantly through oxygenation pathways. In contrast, the IFAS process favored the transformation of saturated and reduced precursors, resulting primarily in unsaturated products, with dealkylation identified as the prevalent degradation pathway.
3. The microbial communities in the A/O process were dominated by *Firmicutes* and *Proteobacteria*, while the IFAS process was enriched with *Chloroflexota* and *Proteobacteria*. The difference in microbial community composition and the EPS content could explain the differences in the removal and transformation between the two processes.

## Supporting information

Supplemental informations

