## Supplemental informations for "Coupling Anaerobic–Oxic and Integrated Fixed–Activated Sludge Processes for Removal and Transformation of Refractory Organic Matter in Livestock and Poultry Digestate": SI_Coupling Anaerobic–Oxic and Integrated Fixed–Activated Sludge Processes for Removal and Transformation of Refractory Organic Matter in Livestock and Poultry Digestate.pdf

Type of contribution: ***Research Article***

* Corresponding authors.

***Table S1 Variations of organic matter at molecular level***

| Sample | FN | MW | $DBE_{wa}$ | $O/C_{wa}$ | $H/C_{wa}$ | $NOSC_{wa}$ | $MLB_L$ | $AI_{mod}$ |
|---|---|---|---|---|---|---|---|---|
| $A/O_{inf}$ | 6110 | 366.66 | 6.11 | 0.42 | 1.44 | -0.35 | 0.35 | 0.07 |
| A zone | 6381 | 372.78 | 6.36 | 0.43 | 1.42 | -0.29 | 0.33 | 0.07 |
| O zone | 6435 | 372.30 | 6.37 | 0.43 | 1.42 | -0.28 | 0.33 | 0.08 |
| $A/O_{eff}$ | 6379 | 373.83 | 6.42 | 0.42 | 1.41 | -0.28 | 0.33 | 0.08 |
| $IFAS_{inf}$ | 6068 | 369.00 | 6.39 | 0.43 | 1.40 | -0.26 | 0.32 | 0.08 |
| IFAS 2 | 6038 | 366.36 | 6.85 | 0.44 | 1.34 | -0.18 | 0.26 | 0.13 |
| IFAS 4 | 5672 | 362.99 | 6.87 | 0.45 | 1.33 | -0.14 | 0.24 | 0.13 |
| $IFAS_{eff}$ | 6007 | 364.56 | 6.96 | 0.44 | 1.33 | -0.16 | 0.24 | 0.14 |

***Text 1 Reactor set-up and operation***

The coupled system comprised five main components: an A/O reactor, a primary settling tank, a regulating tank, an IFAS reactor, and a secondary settling tank. All units were constructed from Plexiglas. The A/O reactor (50 cm × 30 cm × 33.5 cm; 50 L total volume) was divided into four chambers (12.5 L each). The first two chambers were equipped with mechanical mixers to maintain anaerobic conditions, while the last two were aerated using air pumps installed at the bottom to supply oxygen for microbial activity. The primary settling tank (30 L) was a vertical-flow sedimentation unit with a sludge hopper for gravity-based sludge separation and collection. The regulating tank (80 L) served as a buffer between the A/O and IFAS reactors, collecting the A/O effluent and delivering it to the IFAS unit. The IFAS reactor shared the same length and width as the A/O reactor (50 cm × 30 cm) but had a greater depth (73.5 cm), yielding a total volume of 110 L. It was divided into four chambers (27.5 L each), each fitted with mechanical mixers and air pump. Stainless-steel racks were installed in each chamber to hold biofilm carriers, which consisted of 25 mm plastic balls and 20 mm polyurethane sponges. The secondary settling tank (40 L), similar in structure to the primary one, was also a vertical-flow sedimentation unit.

Undiluted livestock and poultry digestate flowed through the A/O reactor in an “S”-shaped pattern to enhance mixing and contact. A portion of the sludge settled in the primary tank was recycled to the anaerobic zone of the A/O reactor at a return ratio of 250%. Effluent from the regulating tank was supplied to the IFAS reactor. To investigate the effect of influent dilution, the proportion of regulating tank effluent gradually increased from 10% to 100% in 10% increments. The temperatures of the A/O and IFAS reactors were maintained at 32 ± 1°C, while other components operated at ambient temperature. The hydraulic retention time (HRT) ranged from 0.5 to 3.0 days for the A/O reactor and from 1.0 to 4.5 days for the IFAS reactor, depending on operational conditions.

*Text S2 2D-COS analysis*

The 2D-COS data processing is to calculate the synchronous and asynchronous correlation intensities at each pair of variables using measurement responses at corresponding pair of variables (Chen et al., 2015a). Specifically, all input data can be represented as a function of an independent variable ($p$) and an external variable $t$ at $J$ evenly spaced points:

$$y(p, t_j), j = 1,2, \dots, J \tag{1}$$

where $p$ represents a fluorescence component in the $F_{max}$ data. The dynamic spectrum is defined as follows (Hur et al., 2011):

$$\tilde{y}(p,t) = \begin{cases} y(p,t) - \bar{y}(p), & T_{min} \leq 0 \leq T_{max} \\ 0, & otherwise \end{cases} \tag{2}$$

$$\bar{y}(p) = \frac{1}{T_{max} - T_{min}} \int_{T_{min}}^{T_{max}} y(p,t)\, dt \tag{3}$$

The Hilbert-Noda discrete transform is used to compute a pair of synchronous ($\Phi$) and asychronous ($\Psi$) correlation intensities (Hur et al., 2011):

$$\Phi(p_1, p_2) = \frac{1}{m-1} \sum_{j=1}^{m} \tilde{y}(p_1, t_j) \tilde{y}(p_2, t_j) \tag{4}$$

$$\Psi(p_1, p_2) = \frac{1}{m-1} \sum_{j=1}^{m} \tilde{y}(p_1, t_j) \sum_{k=1}^{m} N_{jk} \tilde{y}(p_2, t_k) \tag{5}$$

$$N_{jk} = \begin{cases} 0, & if\ j = k \\ \frac{1}{\pi(k-j)}, & otherwise \end{cases} \tag{6}$$

The number of dimensions of the output $\Phi$ and $\Psi$ were determined by the dimension of the variable $p$ of input data in Eq.(1). In the EEM-PARAFAC-COS analysis, the final $\Phi$ and $\Psi$ maps were discrete because the variable of $F_{max}$ (i.e., the number of components) was discrete. The Rayleigh scattering and Raman scattering in EEM data, which were non-informative and would impact the COS computation, were removed prior to analysis by using Matlab and the drEEM toolbox, and the inner-filter effect on EEM data was corrected using the drEEM tool box (Murphy et al., 2013).

***Text S3 Solid-phase extraction***

Agilent Bond Elut-PPL cartridges (500 mg, 6 mL) were pre-conditioned with methanol prior to use. Water samples were acidified to pH 2 using hydrochloric acid (HCl) and passed through the pre-rinsed cartridges to retain dissolved organic matter (DOM). The cartridges were subsequently washed with 0.01 M HCl and dried under a nitrogen stream. DOM was then eluted with methanol, and the eluate was evaporated to dryness under gentle nitrogen flow (Chen et al., 2015b).

***Text S4 Attributes of DOM molecules and samples***

(1) Molecular relative intensity ($M_i$):

$$M_i = \frac{I_i}{\sum I_i}$$

(2) Average molecular weight ($MW_{wa}$):

$$MW_{wa} = \sum (MW_i \times M_i)$$

(3) Double bond equivalent (DBE):

$$DBE = \frac{2 \times C + N + P - H + 2}{2}$$

(4) Average DBE ($DBE_{wa}$):

$$DBE_{wa} = \sum (DBE_i \times M_i)$$

(5) DBE−O per carbon ($(DBE - O)/C$):

$$(DBE - O)/C = \frac{(2 \times C + N + P - H + 2)/2 - O}{\mathrm{C}}$$

(6) Nominal oxidation state of carbon (NOSC):

$$NOSC = 4 - \frac{4 \times C + H - 2 \times O - 3 \times N - 2 \times S + 5 \times P}{\mathrm{C}}$$

(7) Average NOSC ($NOSC_{wa}$):

$$NOSC_{wa} = \sum (NOSC_i \times M_i)$$

(8) Average ratio of oxygen to carbon ($O/C_{wa}$):

$$O/C_{wa} = \sum ((O/C)_i \times M_i)$$

(9) Average ratio of hydrogen to carbon ($H/C_{wa}$):

$$H/C_{wa} = \sum ((H/C)_i \times M_i)$$

(10) Modified aromaticity index ($AI_{mod}$):

$$AI_{mod} = \frac{1 + C - 0.5 \times O - S - 0.5 \times (N + P + H)}{\mathrm{C} - 0.5 \times O - N - S - P}$$

(11) Molecular lability index ($MLB_L$):

$$MLB_L = \frac{N_{H/C>1.5}}{N_t}$$

Here, $M_i$, $I_i$, $MW_i$, $DBE_i$, $NOSC_i$, $O/C_i$ and $H/C_i$ were the relative magnitude, the detected peak magnitude, the molecular weight, DBE, NOSC, O/C and H/C for each formula; C, H, O, N, and S were the number of carbons, hydrogen, oxygen, nitrogen, and sulfur atoms; $N_{H/C>1.5}$was the number of formula with H/C ratio > 1.5, and $N_t$ was the total number of formulas in the water samples (Wang et al., 2021).

***Text S5 Mass spectrometry network analysis***

To further elucidate reaction pathways, a reaction pathway analysis was conducted using a well-established R script to generate mass difference networks (Lin et al., 2024). The process was as follows:

(1) The molecular weight of each compound was precisely calculated to six decimal places using atomic masses approved by the International Union of Pure and Applied Chemistry (IUPAC) for elements (C, H, O, N, S, and P).

(2) Coexisting molecular formulas were identified by comparing samples before and after treatment.

(3) The relative peak intensity abundance was evaluated for coexisting formulas using the ratio fc (peak intensity after treatment / peak intensity before treatment). Precursors were compounds present in the initial sample but absent in the subsequent sample, as well as compounds with an $fc < 0.5$; Resistants included compounds found in both pre- and post-treatment samples, with $0.5 \leqslant fc \leqslant 2.0$; and products consisted of compounds that were absent in the initial sample but emerged in the subsequent sample, including those with $fc > 2.0$.

(4) Linkage analysis was performed to identify potential precursor–product pairs based on exact mass differences.

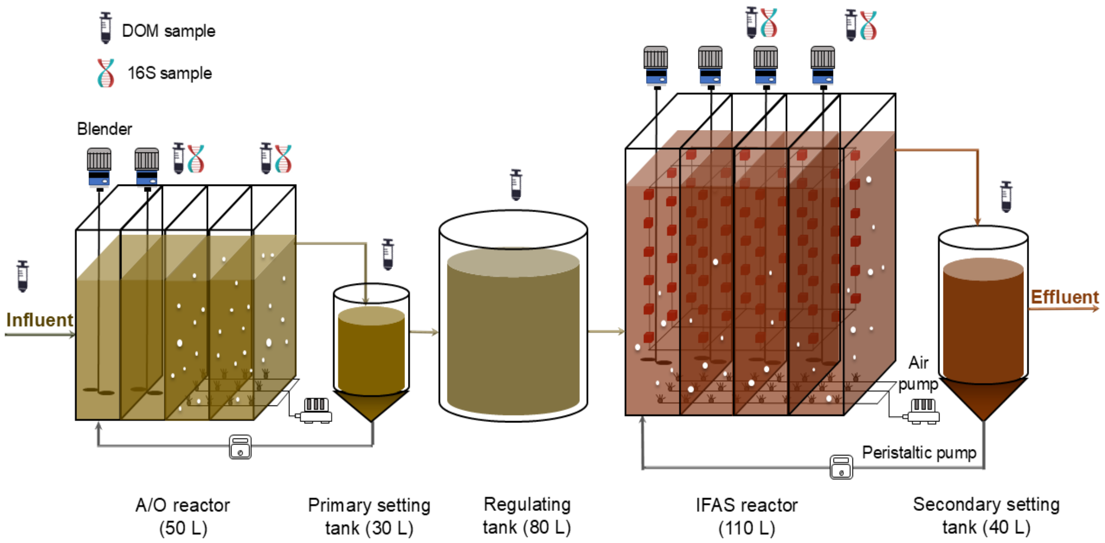


***Fig. S1. Schematic diagram of AO-IFAS process***

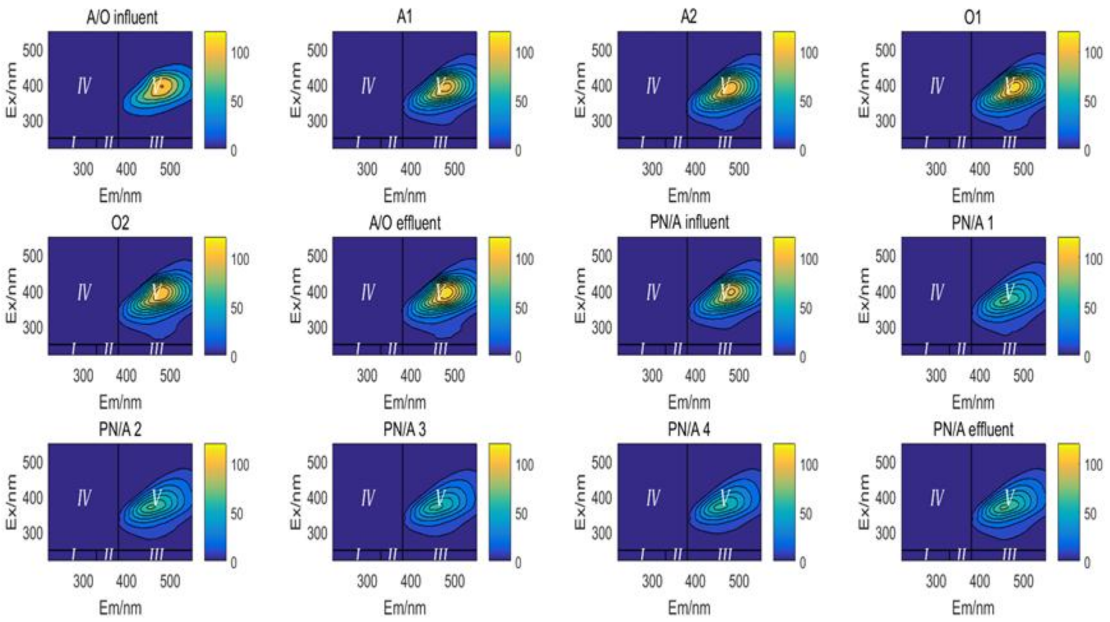


***Fig. S2. Variation of fluorescence spectra along the treatment***

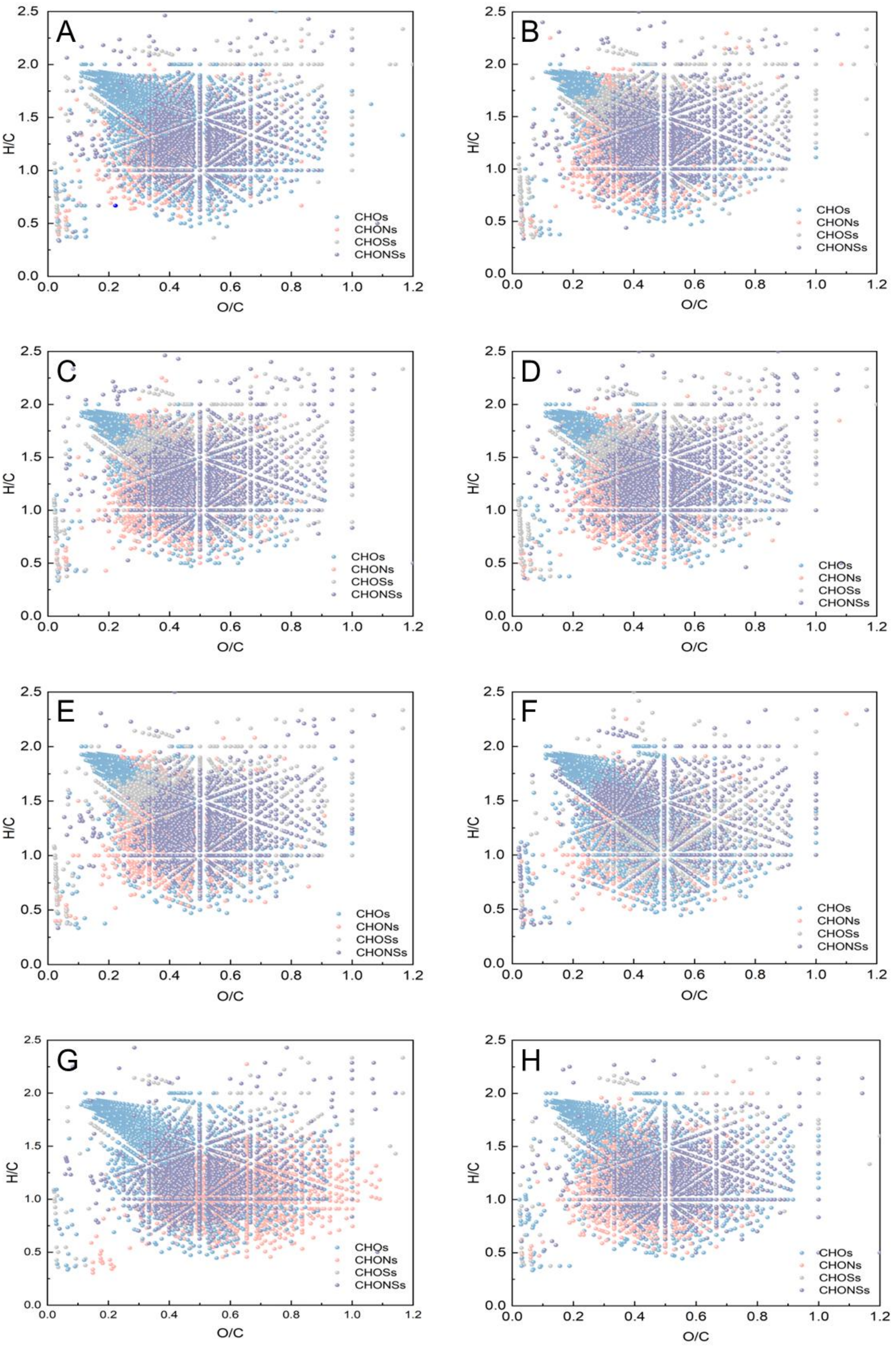


***Fig. S3. Van Krevelen diagrams for DOM formulas. (a) A/O$_{inf}$, (b)A zone, (c) O zone, (d)AO$_{eff}$, (e)IFAS$_{inf}$, (f) IFAS 2, (g)IFAS 4 and (h)IFAS$_{eff}$.***

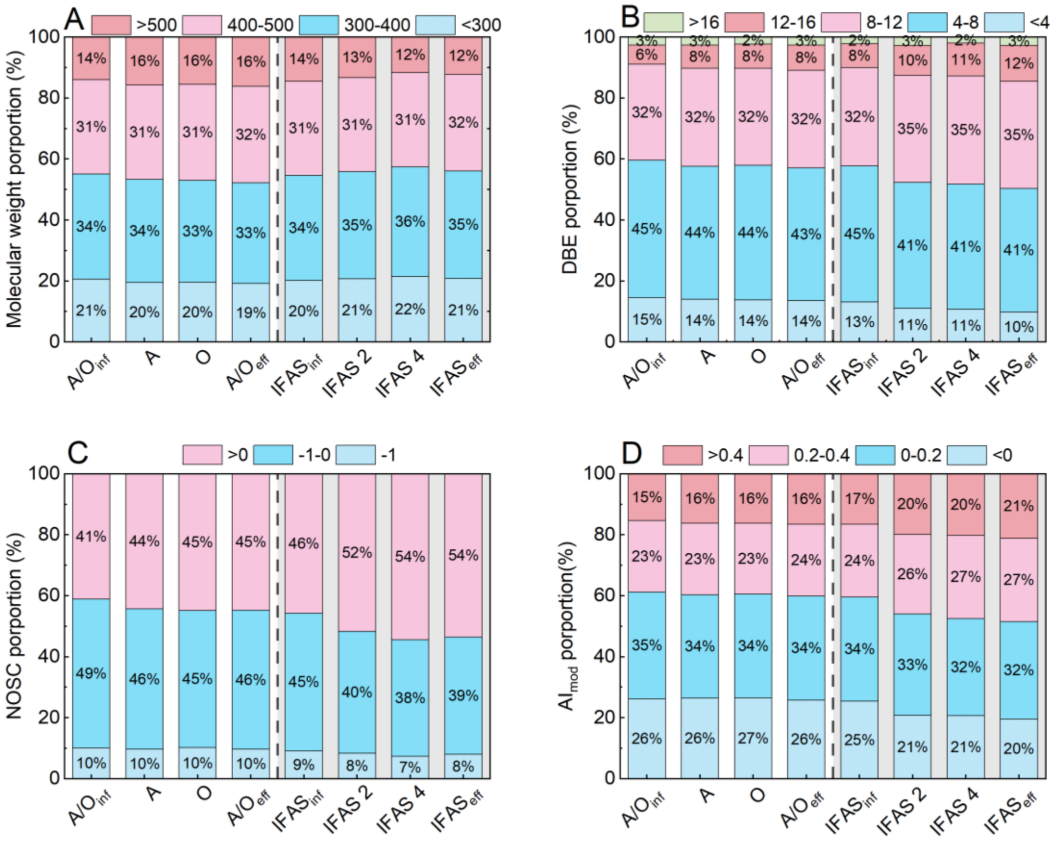


***Figure S4. Distribution of (A) molecular weight, (B) DBE, (C) NOSC and (D) AImod of the refractory organic matter in the A/O and IFAS processes.***

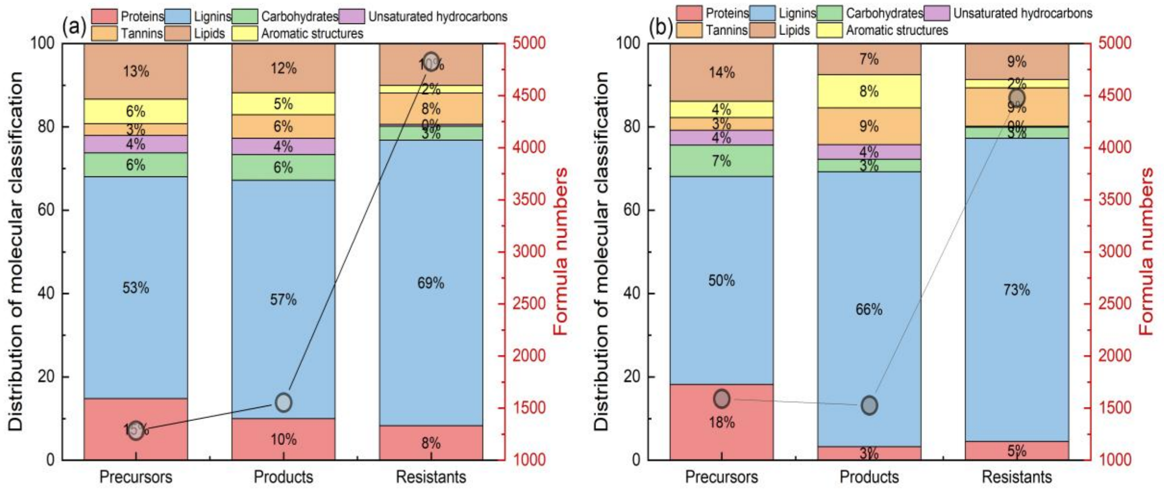


*Fig. S5. Classification of precursors, products, and resistants in (A) A/O reactor and (B) IFAS reactor and corresponding molecular numbers*

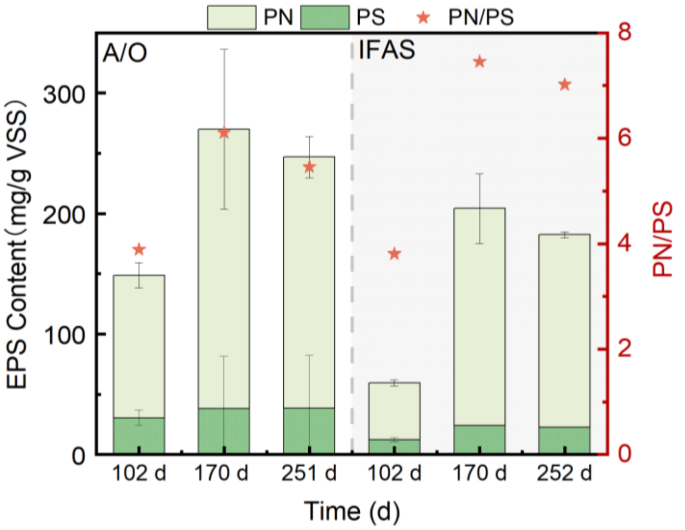


*Fig. S6. Long-term variation of PN, PS and PN/PS ratio.*
